# Simulated green turtle grazing reduces seagrass productivity and alters benthic community structure while triggering further disturbance by feeding stingrays

**DOI:** 10.1101/2022.09.27.509764

**Authors:** Abigail Libbin Cannon, Michael G Hynes, Mackenzie Brandt, Christian Wold, Aaron O’Dea, Andrew Altieri, Jennifer E Smith

## Abstract

While green turtles (*Chelonia mydas*) were once abundant throughout the Caribbean, overexploitation has dramatically reduced their numbers. We conducted a 168-day simulated grazing experiment to determine how loss of this once-abundant mega-herbivore could have affected the productivity and community composition of *Thalassia testudinum-dominated* seagrass beds in Bocas del Toro, Panama. Simulated grazing reduced both percent cover and productivity of *T. testudinum*. High runoff and local pollution from industrial farming may limit light availability and reduce seagrass photosynthetic performance to replace biomass lost to simulated grazing. Other seagrass species and algae failed to colonize space opened by reductions in *T. testudinum* percent cover. Many plots subjected to simulated grazing were also bioturbated by stingrays. Relevance of these findings to balancing sea turtle and seagrass conservation efforts are discussed.

## INTRODUCTION

The historical exploitation of green turtles (*Chelonia mydas*) is estimated to have reduced their abundance in the Caribbean to less than 1% of their pre-Colombian levels (McClenachan et al., 2006). The resultant reduction in grazing pressure has altered benthic community structure (Hearne et al., 2019; Molina Hernández & van Tussenbroek, 2014) and productivity (Holzer & McGlathery, 2016; Molina Hernández & van Tussenbroek, 2014; Moran & Bjorndal, 2005; Williams, 1988; Zieman et al., 1984) of Caribbean seagrasses. Studies of seagrass beds throughout the Caribbean began after *C. mydas* populations had already been substantially reduced, leaving us with a “shifted baseline” of Caribbean seagrass ecology (Jackson et al., 2001; McClenachan et al., 2006) where most studies of seagrass beds have taken place under conditions of artificially reduced megafaunal herbivory. Further, no previous studies have taken place in the Panama-Costa Rica runoff region where Bocas del Toro, Panama is located (Chollett et al., 2012). This region experiences consistently low salinity due to runoff and river discharge caused by high rainfall (Chollett et al., 2012), which can influence which species are present in seagrass beds (Biber & Irlandi, 2006). Sediment-laden runoff can also further stress marine plants by reducing water clarity and light availability (D’Croz et al., 2005), which can negatively affect the ability of seagrass to respond to disturbances (Eklöf et al., 2009).

The Panama-Costa Rica runoff region may also be especially relevant to green turtle conservation due to its close proximity to Tortuguero, Costa Rica, the largest remaining green turtle rookery in the Atlantic Ocean (Troëng & Rankin, 2005). Turtles that nest in Tortuguero and feed nearby may be able to put more energy towards reproduction than turtles that have to travel greater distances between feeding and nesting sites (Troëng et al., 2005). Efforts to help Caribbean green turtles recover would benefit from understanding the likely impact of grazing on the seagrass beds that may sustain some of the most fecund females.

Fourqurean et al. (2010) proposed that seagrass productivity should respond favorably to grazing when the energetic benefits of reduced self-shading are greater than the energetic costs of having to replace lost biomass. Holzer and McGlathery (2016) suggested that grazing should only enhance seagrass production when seagrasses are not limited by phosphorous and did not discuss environmental light limitation. These hypotheses and observations can be combined to predict that seagrasses should benefit from grazing only when they are not nutrient limited, and experience light limitation caused by self-shading rather than turbid water. That most studies of real or simulated grazing have observed reduced productivity (Fourqurean et al., 2010; Greenway, 1974; Molina Hernández & van Tussenbroek, 2014; Williams, 1988; Zieman et al., 1984) (but see Moran & Bjorndal, 2005) suggests that the cost of replacing leaf tissue lost to grazing is usually greater than the benefit of reduced self-shading.

Green turtles have been documented to repeatedly re-grazed the same seagrass patches (Bjorndal, 1980; Fourqurean et al., 2010; Molina Hernández & van Tussenbroek, 2014) and this may be nutritionally advantageous. Nutrient content of seagrass tissue decreases as it ages (Bjorndal, 1980) and nutrients are translocated away from senescing tissue (Hemminga et al., 1999). Grazed blades have a higher young tissue to old tissue ratio than ungrazed ones giving them higher concentrations of nitrogen and phosphorous per unit of biomass (Moran & Bjorndal, 2007). From the seagrass’s perspective simulated herbivory reduces rhizome non-structural carbohydrate content in *T. testudinum* (Fourqurean et al., 2010) as carbohydrates are translocated to fuel leaf growth. This might logically be expected to lead to higher ^13^C content and less negative *δ*^13^C value of seagrass leaves reflecting leaf growth fueled by starch reserves rather than recently fixed carbohydrates (Maunoury-Danger et al., 2010). Changes in *δ*^13^C must, however, be interpreted with caution, because increased light availability may also lead to less negative values (Durako & Hall, 1992), particularly if grazing leads to reduced self-shading, although this is most likely to occur in high shoot density meadows (Enríquez & Pantoja-Reyes, 2005).

In addition to changing the physiology of individual seagrass plants, grazing can also change seagrass community structure. Studies of the impacts of sea turtle grazing in both the Caribbean (Molina Hernández & van Tussenbroek, 2014) and the Indo-Pacific (Heithaus et al., 2014; Kelkar et al., 2013) have shown a tendency of intensely grazed communities to show decreased abundance of late-successional species in the genus *Thalassia* and increased abundance of rhizophytic green algae and smaller, early successional seagrass genera such as *Syringodium* or *Halophila* (but see Hearne et al., 2019).

Grazing by *C. mydas* on seagrass may also alter the behavior of bioturbators. For example, Williams (1988) observed stingray feeding pits in a seagrass bed subjected to heavy disturbance by *C. mydas* (and boat anchors) in the US Virgin Islands. Fourqurean et al. (2010) also observed enhanced bioturbation of seagrass beds grazed *C. mydas* in Bermuda but did not identify the organisms responsible. Stingrays, however, are not thought to be able to disturb continuous beds of ungrazed *T. testudinum* (Valentine et al., 1994). Because stingray disturbance may interfere with the recovery of grazed seagrass patches and, more research is needed on how stingrays respond to disturbances in seagrass beds.

Finally, as the importance of seagrass beds as carbon sinks is increasingly recognized (Fourqurean et al., 2012) it is important to understand how megafaunal grazing affects this process. Reductions in seagrass canopy height caused by sea turtle grazing may lead to reduced particle trapping or greater erosion of high-carbon surface sediments and thus lower carbon storage (Heithaus et al., 2014; Johnson et al., 2019). Johnson et al. (2019), however found simulated *C. mydas* grazing to have no effect on surface sediment carbon content in the Cayman Islands.

In this study we explore how simulated *C. mydas* grazing affected experimental plots in *Thalassia testudinum*-dominated seagrass beds in Bocas del Toro, Western Panama. *C. mydas* were historically abundant in Bocas del Toro (Fig. 1) (Wake et al., 2013) but are now extremely rare in the region (Meylan et al., 2013) and protected by Panamanian and international law (Ankersen et al., 2015; Ruiz et al., 2007). While enforcement of these protections is variable (Ruiz et al., 2007), the number of nests laid per year is increasing at Tortuguero, Costa Rica (Troёng & Rankin, 2005). Tortuguero is the nesting beach for most of Bocas del Toro’s green turtles (Meylan et al., 2013) and the Atlantic Ocean’s largest remaining sea turtle rookery (Troёng & Rankin, 2005). An increase in sea turtles is likely to lead to increased grazing pressure on Bocas del Toro’s seagrass beds. We conducted a 168-day (6 lunar month) simulated grazing experiment to assess how varying levels of sea turtle grazing pressure affects benthic community structure, seagrass productivity, seagrass morphometrics, seagrass leaf nutrient content and *δ*^13^C, seagrass rhizome non-structural carbohydrate content, and sediment carbon content. The results can provide insight into the Caribbean’s past and future under successful regeneration of *C. mydas* populations in highly runoff-influenced tropical marine ecosystems.

**FIG 1.**
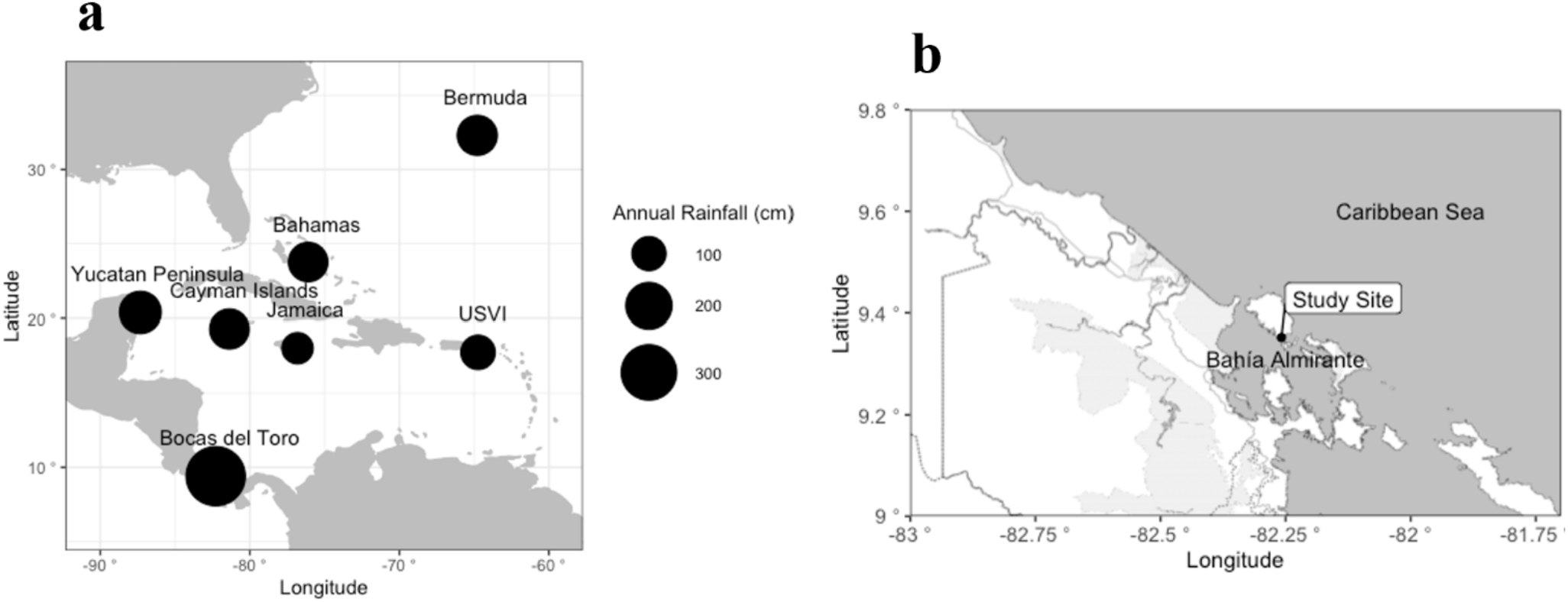
Average annual rainfall in Bocas del Toro, the site of this experiment, as well as at the sites of previous sea turtle grazing studies in the Caribbean and the location of our study within Bocas del Toro.

## MATERIALS & METHODS

### SETTING

We focused our study in the Bocas del Toro region of western Panama (Fig. 1a). Wake et al. (2013) found evidence that green turtles were an important food resource for pre-Colombian indigenous populations dating back to at least AD 800 and Europeans knew this region was important green turtle habitat since at least the 17^th^ century (Meylan et al., 2013). Seagrass beds in this region are typically dominated by *Thalassia testudinum* (turtle grass) (Guzmán et al., 2005), which is the preferred food source of *Chelonia mydas* in the Caribbean (Bjorndal, 1980; Bjorndal, 1985; Mortimer, 1981). The ecosystems of Bocas del Toro in particular, and the Southwestern Caribbean in general, have been less thoroughly studied than those of the Antilles, the Bahamas, The Cayman Islands, or the Yucatán Peninsula (Collin, 2005).

An important contrast between Bocas del Toro and the sites of previous sea turtle grazing experiments in the Caribbean is that Bocas del Toro receives nearly 3.5 m of rain a year (López-Calderón et al., 2013), making it a significantly wetter climate than the sites of previous sea turtle grazing experiments in the Caribbean (Fig. 1a). Historic dominance of sediment-tolerant (Rogers, 1990) corals in the genus *Porites* (Aronson et al., 2014; Collin, 2005; Guzmán & Guevara, 1998; Seemann et al., 2014) also suggests naturally higher rates of sedimentation than at other sites of previous Caribbean sea turtle grazing studies (Guzmán & Guevara, 1998).

Bocas del Toro’s marine environment has shown signs of anthropogenic stress since the 1950s (Cramer et al., 2015) and possibly earlier (Cramer et al., 2017). Deforestation (Aronson et al., 2014; Guzmán & Guevara, 1998; Seemann et al., 2014) and the establishment of banana plantations (Cramer, 2013; Cramer et al., 2015; Guzmán & Guevara, 1998) have increased erosion and sedimentation. Mangrove clearing has additionally reduced natural sediment trapping (Granek & Ruttenberg, 2008). This increase in sediment pollution has contributed to the replacement of *Porites spp*. by *Agaricia spp*. on coral reefs in the region (Aronson et al., 2014). Nutrients from fertilizer in runoff (Cramer, 2013; Seemann et al., 2014) and untreated sewage from an increasing human population (Cramer, 2013) also further contribute to eutrophication.

Seagrass beds in Bocas del Toro, however, have fared better than coral reefs. Their invertebrate communities are less altered (Fredston-Hermann et al., 2013) and monitored seagrass biomass and leaf area per plot actually increased between 1999 and 2010, although without a corresponding increase in production (López-Calderón et al., 2013). No consistent intra-annual variation was observed in these variables (López-Calderón et al., 2013) suggesting that seasonality is not a major factor in bocatoreño seagrass beds.

To explore the impacts of reduced grazing through the functional extinction of *C. mydas* on the productivity and community composition of seagrass beds we conducted our study at 3-4 m depth in a *Thalassia testudinum* dominated seagrass bed on Isla Colón, Bocas del Toro, Panama (9.3526° N, 82.2583° W) (Fig. 1b). The studied area is located in an enclosed lagoon within the larger Bahía Almirante and is surrounded by mangrove forest on three sides. While *T. testudinum* was the only seagrass species present in the experimental area, rhizophytic algae in the genera *Caulerpa* and *Halimeda* were present in small numbers in and around experimental plots.

### EXPERIMENTAL MANIPULATIONS

We conducted an herbivore manipulation experiment with three levels of disturbance frequency: high frequency clipping (HFC), low frequency clipping (LFC), and unclipped (UC). Five 2-m by 2-m plots were established for each treatment and plot boundaries were marked with floats attached to PVC stakes at each corner in water 3 to 4 meters deep. Green turtle grazing was simulated in the HFC and LFC plots by clipping all seagrass leaves with scissors to within 5 cm of the sediment surface. HFC plots were clipped once every two weeks, LFC plots were clipped once every four weeks, and UC plots were never clipped. To avoid edge effects, a 0.5- to 1-m buffer zone was established around each plot and subjected to the same treatment as the plot itself, but data were not collected in buffer zones. At least 1 m of space was maintained between buffer zones of different plots, but plots were not trenched. Plots showed no signs of stingray feeding pits at the start of the experiment, although rays were frequently seen in the study area. Within the 2-m by 2-m clipped plots, cut blades were collected and removed from the experimental area. Seagrass blades from buffer zones were allowed to float to the surface where they were dispersed by currents. The experiment ran from September 21^st^, 2015, to March 1^st^, 2016, during which time HFC plots were clipped 12 times and LFC plots were clipped 6 times. *C. mydas* has been extensively documented in the Caribbean to re-graze particular patches of seagrass (Bjorndal, 1980; Fourqurean et al., 2010; Molina Hernández & van Tussenbroek, 2014) and our disturbance frequency was within the range applied in previous experiments (Holzer & McGlathery, 2016).

### SEAGRASS PRODUCTIVITY AND MORPHOMETRICS

Seagrass productivity in experimental treatments was assessed using the shoot marking technique described by Zieman (1974) where shoots were pierced near the green/white interface (see CARICOMP, 2001 Fig. 4a) that exists where *T. testudinum* leaves first emerge from the bundle sheath. Shoots were marked on 8 separate occasions between October 6^th^, 2015, and February 23^rd^, 2016, and harvested 7 days after marking (Fig 2). After shoots were harvested and all growth above the marking scar was treated as old growth while all growth below the marking scar and all leaves without scars were treated as new growth. New growth and old growth were separated by cutting the grass blade perpendicular to the marking scar with a razor blade and new growth and old growth were dried to constant mass at 60°C. New growth dry masses were multiplied by shoot density to determine production per square meter of grass. Shoot density was measured by counting all the shoots in two haphazardly placed 25 × 25 cm quadrats within the experimental plot on the day shoots were marked and taking the mean of the two values. Unfortunately, no density or productivity data was taken prior to experimental manipulations. Maximum leaf length and leaf width were also measured for all collected shoots. Unfortunately, the first set of new growth data was also not usable, because it was insufficiently dried.

**FIG 2.**
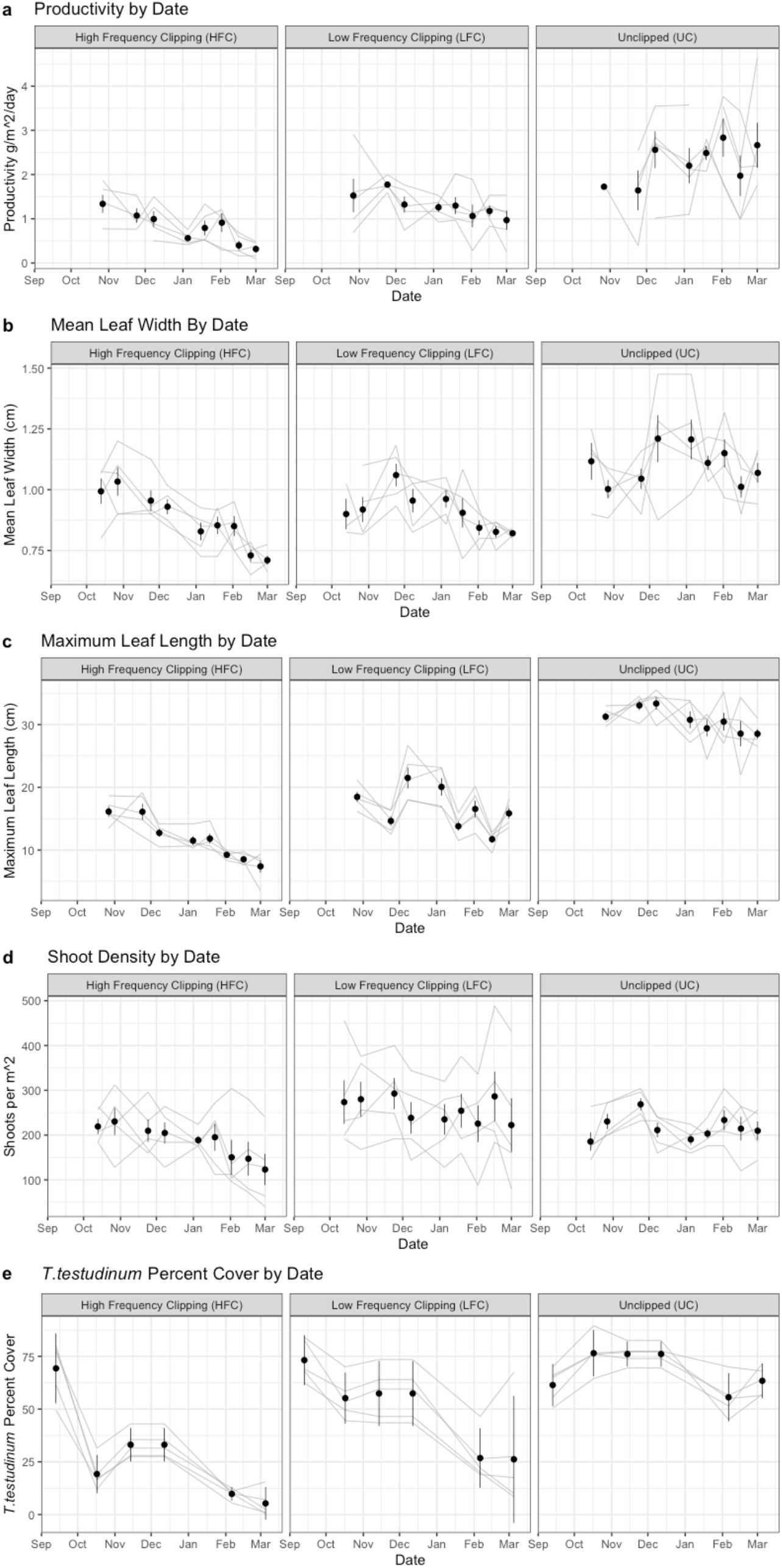
*T. testudinum* production, mean leaf width, maximum leaf length, shoot density, and percent cover over the course of the experiment. Dots indicate means and error bars indicate 95% credible intervals. Lines correspond to individual treatment plots. Breaks in lines indicate times when data for a particular treatment plot was not available.

### SEAGRASS BIOMASS

To estimate initial biomass prior to any experimental manipulations, samples of all above ground macrophyte material was collected in two 25 × 25 cm quadrats placed at opposite corners of the boundaries of each plot and dried to constant weight at 60 °C. To minimize interference of this destructive sampling to the seagrass within plots, the samples were collected from just outside of the plots themselves in the buffer zone. At the end of the experiment biomass was remeasured in the same way except samples were collected in haphazardly placed quadrats from the interiors of plots 20 days after the last clipping treatment of the HFC plots and 48 days after the last clipping treatment of the LFC plots. Belowground biomass was not measured.

### BENTHIC COMMUNITY PERCENT COVER

Percent cover of seagrass, other sessile organisms, and sediment in treatments was quantified prior to the start of the experiment and every 28 days thereafter by photographing all four 1 m^2^ quadrants within each experimental plot with a Canon Powershot G16 camera. This schedule meant that HFC plots were typically photographed 12 days after they were last clipped and LFC plots were typically photographed 16 days after they were last clipped. Images were cropped, white balanced and then analyzed using the image analysis software Photogrid^®^. Photogrid^®^ was programmed to superimpose 50 stratified random points per quadrant on each photo. Organisms under points were identified to the finest possible taxonomic level and points that were over bare sediment rather than organisms were noted as such. Means were taken of the four 1 m^2^ quadrants per plot to determine the percent cover of organisms and sediment for each plot.

### ELEMENTAL COMPOSITION OF SEAGRASS

Elemental composition of seagrass (CNP) was determined using samples of new leaf tissue growth collected on March 1, 2016. Nitrogen and carbon content, and *δ*^13^C was determined using standard combustion in a Costech 4010 autosampler with zero blanks connected to a Thermo Delta XP Mass Spectrometer at Scripps Institution of Oceanography. Phosphorous content of seagrass was determined via flow injection analysis with a Lachat QuikChem 8000 at the Northern Arizona University Environmental Analysis Laboratory. Non-structural carbohydrate analysis of seagrass rhizome tissue collected on March 8^th^, 2016 was conducted using the extraction methods described by Chow and Landhäusser (2004) with subsequent detection of carbohydrate content using the phenol sulfuric acid assay.

### SEDIMENT CARBON CONTENT

To test the impact of simulated grazing on surface sediment carbon content we collected roughly 20 ml of surface sediments in each plot in September 2015 prior to any experimental manipulations and in March 2016 after the end of the experiment. Samples were dried to constant weight at 60°C and were then dry combusted in a Thermo Flash 1112 CHN analyzer at the STRI Soils Lab to determine percent carbon.

### STINGRAY FEEDING

The cumulative number of new stingray feeding pits in each plot were counted by divers throughout the experiment. Feeding pits were identified by their sudden appearance, depth of at least 5cm, and rounded or trench-like shape. New feeding pits were identified by presence in areas that had previously not been “pitted” or having steep sides with no signs of infilling. “Pitted” areas of plots infilled with sediment over time, unless they were “re-pitted”, but typically remained dominated by sediment rather than seagrass. Plots showed no signs of stingray feeding pits prior to the start of the experiment.

### DATA ANALYSIS

Bayes factors comparing the likelihood of models that include fixed effects of date and treatment against the likelihood of a null model including only the random effects of individual experimental plots were compared for seagrass productivity, shoot density, maximum leaf length, leaf width, and leaf area using the generalTestBF function in the ‘BayesFactor’ package (Morey et al., 2021). Seagrass new growth *δ*^13^C, rhizome non-structural carbohydrates, and sediment carbon content were analyzed using an independent variances 1000 iteration one-way ANOVA model in the ‘rjags’ package (Plummer et al., 2018). This model does not require homogenous variance between samples and is robust to deviations from normality (D. Golicher pers comm.). Data were not transformed prior to analysis. The relationship between *T. testudinum* biomass at the end of the experiment and productivity per m^2^ was analyzed using Bayesian robust correlation in the ‘rjags’ package (Plummer et al., 2018) with stan (Stan Development Team, 2018) code developed by Baez-Ortega (2018) based on the work of Bååth (2013). Bayesian logistic regression was conducting using the ‘rjags’ package (Plummer et al., 2018) to calculate the posterior probability distributions of the odds of a plot in different treatments being pitted by sting rays at least once. This was done using a 3 chain, 1000 iteration Markov Chain Monte Carlo model with uninformative priors and a likelihood distribution generated by observations of which plots were pitted. The same package was used to fit a Poisson generalized linear model to the length of time in days between the start of the experiment and the first appearance of a stingray feeding pit in a plot as well as the time in days between any subsequent pitting events in plots that were pitted repeatedly. This was done using a 3 chain, 1000 iteration Markov Chain Monte Carlo model with uninformative priors and a likelihood distribution generated by observations of how long it took for the first or any subsequent pits to appear in repeatedly pitted plots. Whether treatment had any effect on expected number of pits per plot could not be tested statistically due to observations that appearance of feeding pits in many plots accelerated after the first observation suggesting that the occurrence of pits within a given plot were not independent. All analyses were carried out using R (R Core Team, 2019).

## RESULTS

### SEAGRASS PRODUCTIVITY, BIOMASS, MORPHOMETRICS AND PERCENT COVER

In the case of seagrass productivity per unit area, a model including date, treatment, interactions between date and treatment, and individual plot was approximately 47,000 times more likely than a model that included only the random effect of individual plots. It was also over 100 times more likely than the next most likely model, which did not include interaction between date and treatment (Fig 2a). Bayesian robust correlation also revealed a positive relationship (slope 95% credible interval 0.02-0.05) between final seagrass biomass and productivity across treatments (Fig. 3).

**FIG 3.**
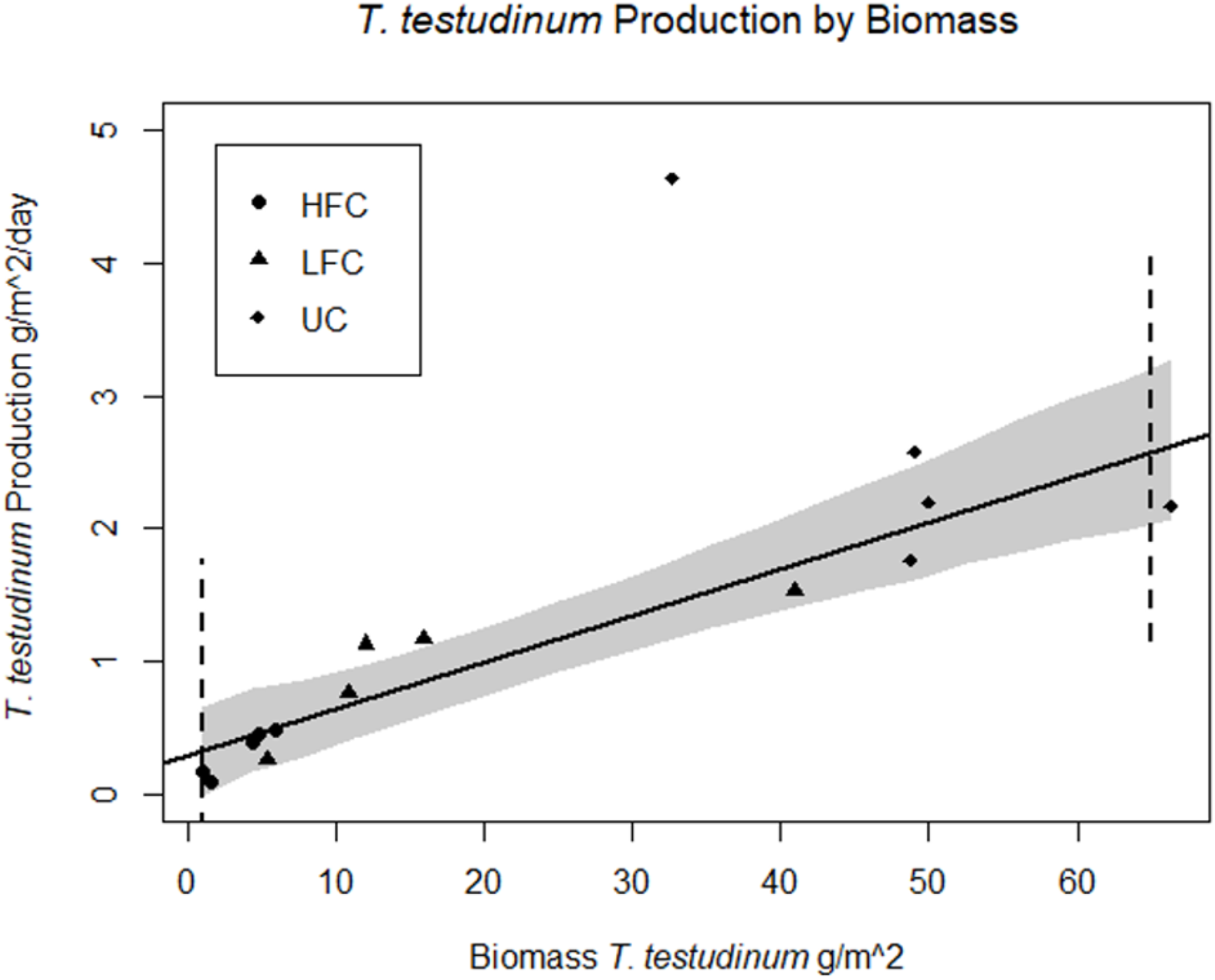
Relationship between aboveground biomass and aerial productivity of *T. testudinum* at the end of the experiment. 95% credible intervals for slope 0.02-0.05

For mean *T*. testudinum leaf width a model including date, treatment, interactions between those fixed effects and random effects of individual plots was over 800 times more likely than the next most likely model which did not include interactions and over 1,600,000,000 times more likely than a model based on plot identity alone (Fig 2b). The case for interactions between date and treatment on maximum leaf length, however, was weaker with a model including treatment, date, interactions between treatment and date, and plot identity only 1.5 times more likely than a model without interactions between date and treatment. A model including all three variables without interactions, however, was still 140,000,000 times more likely than the next most likely model, which included only date and individual plot identity and 2.5×10^16^ times more likely than a model based on plot ID alone (Fig 2c). Shoot density data, however, was noisier with strong individual plot effects. Nonetheless, a model including treatment, date, interactions between treatment and date, and individual plots was 4.7 times more likely than the next most likely model which included only date and plot identity. This model was, however, over 500 times more likely than a model including individual plots as the only variable (Fig 2d). For percent cover of *T. testudinum* a model including date, treatment, interactions between date and treatment was over 1000 times more likely than the next likeliest model, which included all variables without interactions and 2.9 × 10^15^ times as likely as a model including only plot identity (Fig 2e). The green rhizophytic algae and sponges present in small numbers in some experimental plots failed to colonize space made available by *T. testudinum* declines and never averaged more than 1% of benthic cover in any treatment at the end of the experiment.

### ELEMENTAL COMPOSITION OF SEAGRASS AND SEDIMENT

Treatment was not useful for predicting the non-structural carbohydrate content of seagrass rhizomes or for predicting *δ*^13^C of newly grown seagrass leaf tissue (Table 1). Belowground biomass was not measured. Treatment was also not predictive of seagrass new leaf growth carbon or nitrogen content and while phosphorous values in LFC treatments were within the 95% credible interval for UC treatments the measured value for HFC treatments was more than double this range (Table 1). No effect of experimental treatments was observed on sediment carbon content. At the start of the experiment prior to any experimental manipulations 95% credible intervals for sediment carbon content were 7.52-10.20% in UC plots, 8.78-10.30% in LFC plots, and 8.49-10.84% in HFC plots. At the end of the experiment 95% credible intervals for sediment carbon content were 8.85-11.17% in UC plots, 8.72-10.52% in LFC plots, and 8.55-10.61% in HFC plots.

**TABLE 1.**
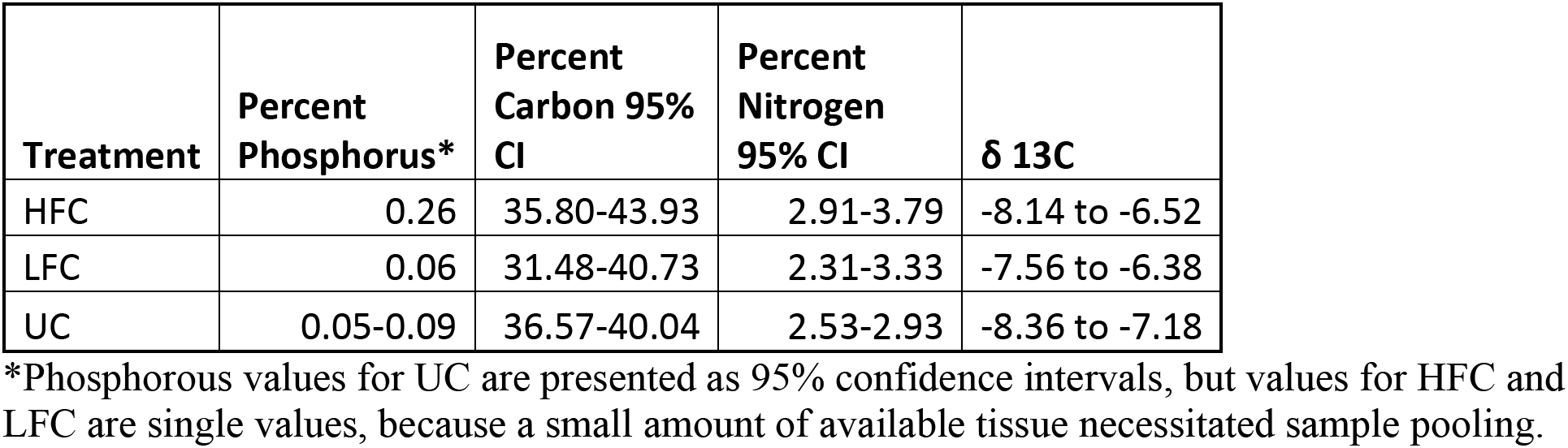
Nutrient content and ratios for new seagrass growth in the high frequency clipping (HFC), low frequency clipping (LFC), and unclipped (UC) treatments.

### STINGRAY FEEDING

The first stingray feeding pit was observed in this experiment in plot HFC2 on December 3^rd^, 2015 (day 73 of the experiment). The first stingray feeding pit observed in a lightly grazed plot was in LFC1 on January 19^th^, 2016 (day 120 of the experiment). No feeding pits were ever observed in UC plots (Table 2). Four out of the six plots where stingray feeding pits were observed would later be re-pitted on different days and time intervals between subsequent “re-pits” were notably shorter than the time interval from the start of the experiment to the first pit in a given treatment (Fig. 4). While southern stingrays (*Dasyatis americana*) were never directly observed while digging in experimental plots, they were the most frequently observed large ray in the area and were observed and photographed digging nearby sand patches within the seagrass bed. Our results show that clipping history (UC vs HFC or LFC) predicted whether a plot was pitted at least once, but the degree of clipping (HFC vs. LFC) did not (Table 3).

**FIG 4.**
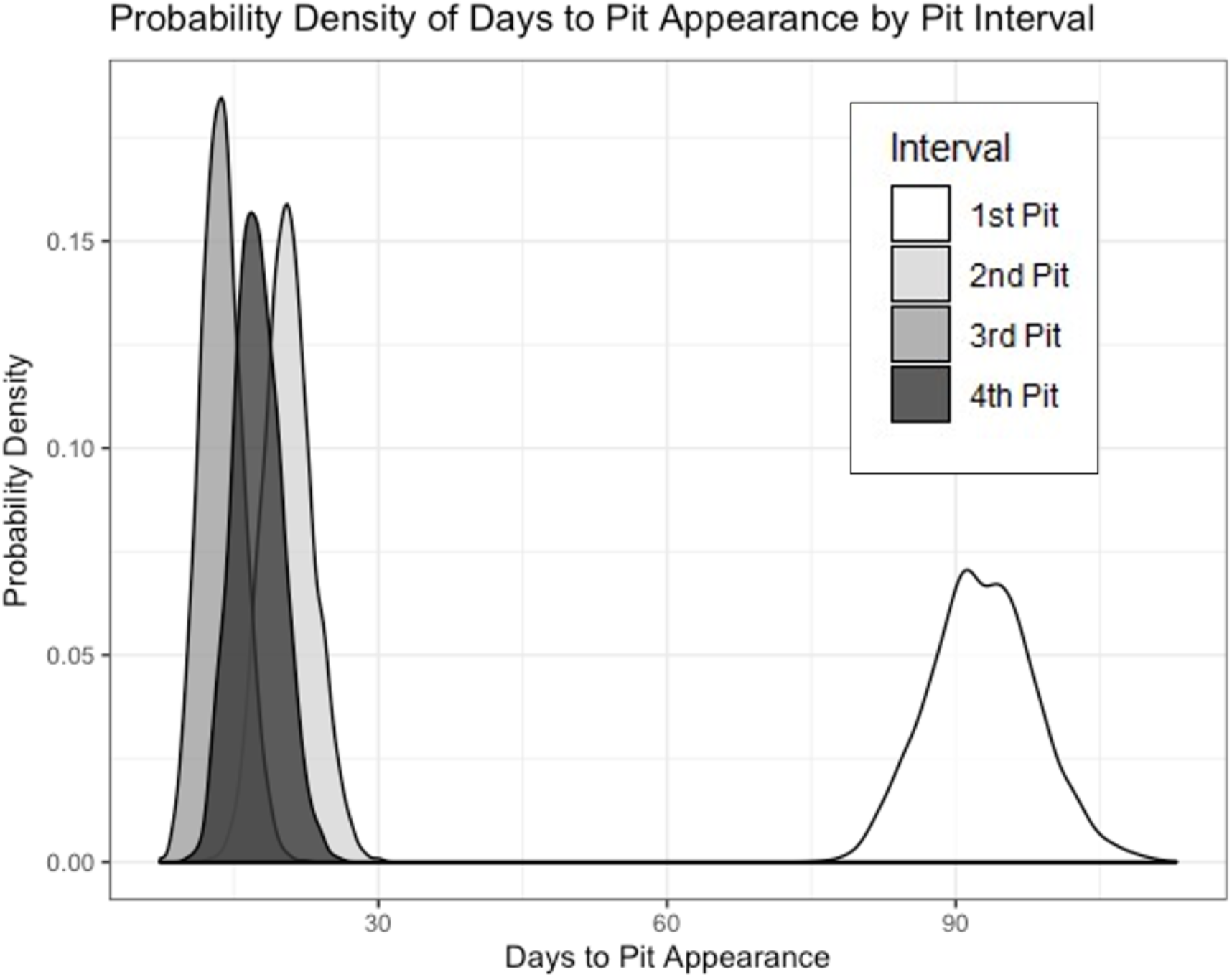
Time in days from start of experiment to appearance of first through fourth stingray feeding pit in a plot. Only pitted plots were used in this analysis. Plots show distribution of values for different treatments generated by an independent variances 1000 iteration one-way ANOVA model in the ‘rjags’ package. Possible values each treatment are depicted on the x-axis. Note: productivity is a continuous variable and while the area under the curve must be equal to one, the y-axis values are not meaningful by themselves and can be greater than or equal to 1.

**TABLE 2.** Total number of pits in each plot by the end of the experiment, date of observation of first feeding pit if applicable, and whether a plot was “re-pitted” if applicable. (UC = Unclipped, LFC = low frequency clipping, HFC = high frequency clipping).

| **Plot** | **Type** | **Number of Ray**<br>**Feeding Pits** | **Days After Start of**<br>**Experiment First Pit**<br>**Observed** | **"Re-Pitted" After**<br>**First Observation** |
| --- | --- | --- | --- | --- |
| HFC1 | HFC | 9 | 86 | Yes |
| HFC2 | HFC | 4 | 73 | Yes |
| HFC3 | HFC | 2 | 100 | No |
| HFC4 | HFC | 0 | NA | NA |
| HFC5 | HFC | 0 | NA | NA |
| LFC1 | LFC | 6 | 120 | Yes |
| LFC2 | LFC | 0 | NA | NA |
| LFC3 | LFC | 1 | 141 | No |
| LFC4 | LFC | 0 | NA | NA |
| LFC5 | LFC | 0 | NA | NA |
| UC1 | UC | 0 | NA | NA |
| UC2 | UC | 0 | NA | NA |
| UC3 | UC | 0 | NA | NA |
| UC4 | UC | 0 | NA | NA |
| UC5 | UC | 0 | NA | NA |

**TABLE 3:** Bayesian medians and minimum and maximum credible intervals for odds of a plot in a given treatment being “pitted” at least once over the course of the experiment. (UC = unclipped, LFC = low frequency clipping, HFC = high frequency clipping.

| Treatment | Min | Median | Max |
| --- | --- | --- | --- |
| UC | 0.00 | 0.00 | 4.17 |
| LFC | 1.43 | 28.12 | 85.06 |
| HFC | 18.79 | 61.11 | 93.86 |

## DISCUSSION

We found that simulating historical grazing by ecologically extinct green sea turtles reduced productivity, biomass, and leaf width in *T. testudinum* seagrass in Bocas del Toro, however no difference in *δ*^13^C content of new leaf tissue or rhizome non-structural carbohydrates was detected. This may be because *T. testudinum* clones can reach over 100 m (van Dijk and van Tussenbroek, 2010) and rhizomes sampled from grazing treatments may have received non-structural carbohydrates translocated from outside our treatment plots. This is particularly relevant since the largest *T. testudinum* clones are found in lagoonal habitats similar to the experimental site (van Dijk & van Tussenbroek, 2010). Undamaged seagrass rhizomes have also been theorized to be sufficient to prevent erosion of surface sediments, especially in environments with low water motion (Johnson et al., 2019), which may explain why no effect of treatment on sediment surface carbon content was detected. This also suggests that increased *C. mydas* grazing does not necessarily affect the capacity of seagrass beds as carbon sinks.

Bocas del Toro experiences a particularly wet climate (Fig. 1a) with extensive rainfall that causes regular reductions in water clarity (D’Croz et al., 2005). These conditions may explain why we found that clipping reduced seagrass productivity even though new leaf tissue was highest in phosphorous in the highly grazed treatment. This also suggests that growth of seagrasses in the experimental site was more strongly limited by light than by nutrients. Mean shoot density in all treatments was less than 300 shoots per meter, while the lowest shoot density at any CARICOMP monitored site in the Caribbean was roughly 400 shoots per meter (van Tussenbroek et al., 2014), reinforcing the idea that any light limitation the plants experienced was the result of environmental light limitation rather than shading by conspecifics, particularly because *T. testudinum* reduces shoot density when subjected to light limitation (Ibarra-Obando et al., 2004). Environmental light limitation may also explain why we did not observe an increase in shoot density in response to simulated grazing as was observed by Holzer and McGlathery (2016) in Bermuda.

The finding that clipping reduced *T. testudinum* productivity agrees with studies of real or simulated sea turtle grazing conducted in the US Virgin Islands (Williams, 1988; Zieman et al., 1984), Bermuda (Fourqurean et al., 2010; Holzer & McGlathery, 2016), Jamaica (Greenway, 1974), and the Yucatan Peninsula (Molina Hernández & van Tussenbroek, 2014), but not with the findings of Moran and Bjorndal (2005) in the Bahamas. The generally detrimental effects of sea turtle grazing on *T. testudinum* leads to questions about how this seagrass species persisted in the Caribbean if *C. mydas* was 300 times more abundant than in the present (McClenachan et al., 2006). One possible explanation is McClenachan et al.’s (2006) estimate of pre-Colombian *C. mydas* abundance is based on females at nesting beaches, not on individuals feeding in seagrass beds and *C. mydas* do not always feed near where they nest. In fact, individuals from the Tortuguero nesting population have been observed feeding as far afield as North Carolina (Bass et al., 2006).

Even allowing for Caribbean *C. mydas* to be subsidized by seagrass beds outside the Caribbean, the finding of an adverse impact of clipping on seagrass productivity in Bocas del Toro is concerning. Bjorndal (1995) estimates that a mature female green turtle needs to eat 100 kg dry weight of *Thalassia testudinum* each year. If we assume that the *T. testudinum* median modeled productivities of 0.97 g m-^2^ day-1 in low frequency clipped (LFC) plots and 0.32 g m-^2^ day-1 in high frequency clipped (HFC) plots are sustainable and that LFC plots could be grazed 12 times over the course of a year, while HFC plots could be grazed 24 times over the course of a year it would take roughly 930 m^2^ of seagrass “pasture” to sustain 1 green turtle at HFC productivity levels or 307 m^2^ of seagrass “pasture” to sustain 1 green turtle at LFC productivity levels. This also means that seagrass meadows in Bocas del Toro could support a maximum of 11 adult female *C. mydas* per hectare at HFC productivity levels or 32 adult female *C. mydas* at LFC productivity levels. This is substantially lower than Bjorndal’s (1995) estimate of 138 adult female *C. mydas* per hectare based on the productivities of clipped seagrass measured by Greenway (1974) in Jamaica. While some of this discrepancy can be accounted for by Bjorndal’s (1995) assumption that *C. mydas* would allow *T. testudinum* biomass to recover to ungrazed levels before re-grazing, an assumption that seems not to describe Caribbean *C. mydas* grazing behavior (Fourqurean et al., 2010; Molina Hernández & van Tussenbroek, 2014) *T. testudinum* productivity also appears to be lower in Bocas del Toro than in Jamaica with or without sea turtle grazing. While Greenway (1974) determined the productivity of ungrazed seagrass in Jamaica to be roughly 3.78 g m-^2^ day-1, the median modeled productivity of unclipped seagrass in our experiment was 2.67 g m-^2^ day-1. Bocas del Toro receives more than three times as much annual rainfall as Jamaica and precipitation in Bocas is less seasonal. We also frequently observed increased water turbidity following intense rain events and this may account for lower productivity in Bocas del Toro than in Jamaica.

Seagrass beds in Bocas del Toro may have lower resilience to grazing than they did in the past as a result of deforestation, coastal development and associated declines in water quality, which has caused phase shifts on coral reefs to more sediment-tolerant coral species (Aronson et al., 2014). Shading slows seagrass recovery from grazing impacts (Eklöf et al., 2009). It is less clear whether the negative response of other Caribbean seagrass communities to sea turtle grazing can be attributed to declining water quality, because they receive less rainfall and are likely less affected by runoff than Bocas del Toro, although deteriorating water quality is a major cause of seagrass losses around the world (Orth et al., 2006). Negative effects of sea turtle grazing on seagrass communities may also be related to sea turtles becoming locally overabundant, even if they are still regionally depleted. Turtle populations may increase in number with the absence of historic levels of shark predation (Heithaus et al., 2014) or shift foraging behavior to more destructive grazing patterns when predator avoidance is no longer necessary (Burkholder et al., 2013).

The inability of other seagrass and algae species to capitalize on declines in *T. testudinum* percent cover, including *Halimeda spp*. and *Caulerpa spp*. already present in the study site, contradicts the findings of Molina Hernández and van Tussenbroek (2014) in the Yucatan Peninsula and Williams (1990) in the US Virgin Islands, but agree with those of Hearne et al. (2019) in the Cayman Islands. These heterogenous responses may be due to differences in grazing pressure by fish or invertebrates (Tribble, 1981) or environmental characteristics being more or less suitable for rhizophytic algae, which grow best in stable “oceanic” environments (Biber & Irlandi, 2006). In our study stingray bioturbation of clipped sites may have interrupted colonization and succession but cannot be the only explanation since not all clipped sites were bioturbated by stingrays and most clipped sites ended the experiment with zero sampled percent cover or biomass of non-T. *testudinum* sessile organisms whether they were bioturbated or not. Grazing is an unlikely explanation, because urchins were rarely seen in the experimental site and green rhizophytic macroalgae are not highly preferred by Caribbean parrotfish (Lobel & Ogden, 1981; Targett et al., 1986). Water clarity appears the most likely explanation for the failure of other organisms to capitalize on declines in *T. testudinum* at our study site, but this would need to be tested systematically by repeating this experiment in less runoff-affected areas and potentially for longer periods of time.

That *T. testudinum* percent cover decreased in clipped plots is not particularly surprising. Clipping obviously leads to shorter seagrass shoots, although our model showed an effect of date as well on treatment on seagrass shoot height, suggesting variability in how much new tissue *T*.t*estudinum* was able to re-grow after clipping. Changes in seagrass shoot width and shoot density over the course of the experiment, however, also suggest that declines in *T. testudinum* percent cover were not exclusively driven by seagrass becoming shorter. The most dramatic declines in *T. testudinum* percent cover across all plots between December and February, which coincided with a period of heavy rainfall and turbid water. Mean percent cover diverges between treatments from February to March with the recovery of unclipped plots, but not clipped ones. This suggests that clipping lowers resilience to other disturbances and this is highly relevant in an area where seagrass beds must frequently contend with reduced light availability due to intensifying sedimentation events (Aronson et al., 2014; Guzmán & Guevara, 1998; Seemann et al., 2014).

The most surprising result of this experiment was the increased likelihood of stingray bioturbation in clipped plots. The only previous mention of the effect of sea turtle grazing on stingray behavior in the literature was by Williams (1988), however it was not clear from that study whether rays were responding to disturbance by grazing or by boat anchor scars. Valentine et al. (1994) noted that rays can dig up disturbed, but not undisturbed continuous areas in *T. testudinum* beds. This is the first study to directly implicate sea turtle grazing in facilitating subsequent disturbance by foraging in seagrass beds. It is not clear, however, whether rays specifically target clipped seagrass patches over other “diggable” habitats, or they simply use disturbed seagrass habitats at about the same frequency they would unvegetated sediments. It is also unclear whether stingrays’ tendency to “re-dig” plots they have previously pitted reflects any enhancement of prey biomass by bioturbation or stingrays simply re-visiting habitats they know to be “diggable” more frequently. Further research into whether sea turtle grazing facilitates stingray feeding, and the extent to which natural or anthropogenic disturbance in general increases stingray “re-disturbance” of seagrass beds is merited.

### IMPLICATIONS FOR *C. MYDAS* CONSERVATION

*C. mydas* does not feed exclusively on seagrass in the Caribbean, although *T. testudinum* provides the greatest contribution to its diet in most studied locations and is the preferred diet along with a few species of red algae (Bjorndal, 1980; Mortimer, 1981). At higher latitudes and in colder water *C. mydas* consumes increasing amounts of animal matter, but this is predicted to become less available as climate change causes water temperatures to increase (Esteban et al., 2020). The results of this experiment and Hearne et al. (2019) also show that declines in *T. testudinum* do not necessarily lead to increases in other macrophyte species or sessile animals. Additionally there is some evidence suggesting that green turtles that feed primarily on algae do not grow as large as those that feed primarily on seagrass (Mortimer, 1995) implying that algae may be an inferior resource. Alternately *C. mydas* may respond to reduced seagrass biomass by migrating away from overgrazed sites to “greener pastures” as was observed by Kelkar et al. (2013) in the Indian Ocean, but declining seagrass health throughout the Caribbean (van Tussenbroek et al., 2014) may limit this strategy. Turtles that have to migrate further between nesting and feeding sites also may not be able to invest as much energy in reproduction as those that do not have to travel as far to meet their nutritional needs (Troëng et al., 2005). All of this indicates that reduced abundance or productivity of *T. testudinum* from deteriorating environmental conditions is likely to have negative impacts on *C. mydas* populations and may limit the ability of modern seagrass beds to sustain turtles at historic levels.

Most sea turtle conservation efforts have so far focused on protecting individual sea turtles, particularly nesting females (Rees et al., 2016) and the success of these efforts can be seen in increasing *C. mydas* nesting at places like Tortuguero, Costa Rica (Troëng & Rankin, 2005). These efforts, however, may not be sufficient to sustainably restore *C. mydas* populations to pre-Colombian levels if modern Caribbean seagrass beds cannot support pre-Colombian grazing pressure. As populations of *C. mydas* increase, organizations dedicated to their conservation may wish to consider the productivity of local *T. testudinum* beds as well as local water clarity when setting *C. mydas* recovery goals. Doing so will ensure their efforts result in a thriving and stable population rather than a population which will unsustainably overgraze its food source. To improve the productivity of seagrass beds in Bocas del Toro, we recommend a reduction in both sediment and nutrient loads contained in terrestrial runoff reaching the ocean (Seemann et al., 2014), which will likely require a combination of reforestation (Gao & Yu, 2017) and reductions in fertilizer use (Seemann et al., 2014). Successful conservation and restoration of *C. mydas* feeding habitat, as well as coral reefs (Aronson et al., 2014; Cramer et al., 2015; Seemann et al., 2014), will also require the conservation and restoration of terrestrial habitats where neither sea turtles nor corals are found.

While much of the Caribbean does not have as wet of a climate as Bocas del Toro, the generally negative response of *T. testudinum* beds to real or simulated *C. mydas* grazing (Fourqurean et al., 2010; Greenway, 1974; Molina Hernández & van Tussenbroek, 2014; Williams, 1988; Zieman et al., 1984) (but see Moran & Bjorndal, 2005) suggests *T. testudinum* today has low resilience to grazing across the Caribbean. Rebuilding resilience will require a cross-ecosystem effort, and this may become increasingly urgent if we wish to restore *C. mydas* populations to pre-Colombian abundances.

## ACKNOWLEDGEMENTS

We are grateful to Levi Lewis, Emily Kelly, Mike Fox, Adi Khen, Bryce Semmens, Brian Stock, and Lynn Waterhouse at Scripps for providing scientific support, especially to Lynn Waterhouse for help with Bayesian statistics. We would also like to thank Davey Kline for the use of your Odyssey Sensor. We are also grateful to the staff of the O’Dea Lab at STRI, Felix Rodriguez in particular, the Castillo family, Plinio Góndola, Deyvis Gonzalez, and Gilberto Murray for providing support in field. Funding for this project was provided by an NSF GRIP Fellowship to ALC and National System of Investigators of SENACYT (Panama) to AO. Frequent flier miles were provided by Chris Cannon.

